## Supplementary file 1 for "A dynamic bactofilin cytoskeleton cooperates with an M23 endopeptidase to control bacterial morphogenesis"

### Supplementary tables

#### Table S1. Strains used in this study.

| Strain | Genotype/description | Construction | Reference/Source |
|---|---|---|---|
| ***H. neptunium*** | | | |
| LE670 | Wild type (*aka* ATCC 15444) | - | Leifson, 1964 |
| EC23 | Δ*HNE_0444* (*bacD*) | In-frame deletion of *bacD* in ATCC 15444 using pEC29 | This study |
| EC28 | Δ*HNE_2629* (*bacA*) | In-frame deletion of *bacA* in ATCC 15444 using pEC32 | This study |
| EC33 | Δ*bacA* Δ*bacD* | In-frame deletion of *bacD* in EC28 using pEC29 | This study |
| EC41 | Δ*bacA* $P_{Cu}$::$P_{Cu}$-*bacA* | Integration of pEC60 in EC28 | This study |
| EC43 | Δ*bacA* $P_{Zn}$::$P_{Zn}$-*bacD* | Integration of pEC61 into EC28 | This study |
| EC60 | Δ*bacA* $P_{Zn}$::$P_{Zn}$-*bacD-venus* | Integration of pEC59 into EC28 | This study |
| EC61 | *bacA*::*bacA-eyfp* | Replacement of *bacA* with *bacA-eyfp* in ATCC 15444 using pEC74 | This study |
| EC67 | *bacD*::*bacD-venus* | Replacement of *bacD* with *bacD-venus* in ATCC 15444 using pEC75 | This study |
| EC68 | *bacA*::*bacA-eyfp bacD*::*bacD-mCherry* | Replacement of *bacD* with *bacD-mCherry* in EC61 using pEC76 | This study |
| EC93 | *HNE_0620*::*eyfp-HNE_0620* (*rodZ*) | Replacement of *rodZ* with *eyfp-rodZ* in ATCC 15444 using pEC129 | This study |
| MO78 | *bacA*::*bacA*$_{F130R}$-*eyfp* | Replacement of *bacA* with *bacA*$_{F130R}$-*eyfp* in ATCC 15444 using pMO93 | This study |
| SP221 | Δ*bacA* Δ*bacD rodZ*::*eyfp-RodZ* | Replacement of *rodZ* with *eyfp-rodZ* in EC33 using pEC129 | This study |
| SP236 | *lmdC*::*lmdC*$_{AA1\text{-}65}$-*HA-lmdC*$_{AA66\text{-}405}$ pdCas9Entry-sgLmdC | Integration of pJH13 in SU34 | This study |
| SP249 | *lmdC*::*lmdC*$_{AA1\text{-}65}$-*HA-lmdC*$_{AA66\text{-}405}$ pdCas9Entry | Integration of pdCas9Entry in SU34 | This study |
| SU34 | *lmdC*::*lmdC*$_{AA1\text{-}65}$-*HA-lmdC*$_{AA66\text{-}405}$ | Replacement of *lmdC* with *lmdC*$_{AA1\text{-}65}$-*HA-lmdC*$_{AA66\text{-}405}$ in ATCC 15444 using pSU23 | This study |
| ***R. rubrum*** | | | |
| S1 | wild type (aka ATCC 11170 or DSM467) | | Molisch, 1907 |
| SP68 | ΔRru_A1868 (*lmdC*) | In-frame deletion of *lmdC* in S1 using pSP81 | This study |
| SP70 | ΔRru_A1867 (*bacA*) | In-frame deletion of *bacA* in S1 using pSP82 | This study |
| SP98 | Δ*lmdC bacA*::*bacA-mNeongreen* | Replacement of *bacA* with *bacA-mNeongreen* in SP68 using pSP119 | This study |
| SP105 | Δ*bacA* $P_{bacA}$-*bacA* | Transformation of SP70 with pSP118 | This study |
| SP109 | *bacA*::*bacA-mCherry* | Replacement of *bacA* with *bacA-mCherry* in S1 using pSP117 | This study |
| SP114 | *bacA*::*bacA-mCherry* $P_{lmdC}$-*lmdC*$_{1\text{-}80}$-*mNeongreen* | Transformation of SP109 with pSP112 | This study |
| SP116 | Δ*bacA* Δ*lmdC* | In-frame deletion of *lmdC* in SP70 using pSP81 | This study |
| SP117 | *bacA*::*bacA-mCherry* Δ*lmdC* | In-frame deletion of *lmdC* in SP109 using pSP130 | This study |
| SP118 | Δ*bacA* Δ*lmdC* $P_{lmdC}$-*lmdC*$_{1\text{-}80}$-*mNeongreen* | Transformation of SP116 with pSP112 | This study |
| SP119 | *bacA*::*bacA-mCherry* Δ*lmdC* $P_{lmdC}$-*lmdC*$_{1\text{-}80}$-*mNeongreen* | Transformation of SP117 with pSP112 | This study |
| SP237 | *bacA*::*bacA-mCherry* Δ*lmdC* $P_{lmdC}$-*lmdC*$_{AA1\text{-}80;\ R25A\ H26A\ L27A}$-*mNeongreen* | Integration of pSP198 in SP117 | This study |
| SP238 | *bacA*::*bacA-mCherry* Δ*lmdC* $P_{lmdC}$-*lmdC*$_{AA1\text{-}80;\ R30A\ S31A}$-*mNeongreen* | Integration of pSP199 in SP117 | This study |
| ***E. coli*** | | | |
| BL21(DE3) | *E. coli* B *dcm ompT hsdS*(rB$^-$ mB$^-$) *gal* | | Invitrogen |
| Rosetta(DE3)pLysS | F$^-$ *ompT hsdS*$_B$($r_B^-$ $m_B^-$) *gal dcm* (DE3) pLysSRARE (Cam$^R$) | - | Merck Millipore |
| TOP10 | F$^-$ *mcr*A Δ(*mrr-hsd*RMS-*mcr*BC) Φ80*lac*ZΔM15-Δ*lac*X74 *rec*A1 *ara*D139 Δ(*ara leu*) 7697 *gal*U *gal*K *rps*L (Str$^R$) *end*A1 *nup*G | | Thermo Fisher Scientific |
| WM3064 | *thrB1004 pro thi rpsL hsdS lacZ*ΔM15 RP4–1360 Δ(*araBAD*)567 Δ*dapA*1341::[*erm pir*(wt)] | - | W. Metcalf (unpublished) |

**Table S2. Backbone plasmids used in this work.**

| Plasmid | Description | Source |
|---|---|---|
| pCCFPC-3 | Integrating plasmid for generation of C-terminal CFP fusions under control of $P_{Cu}$, $Rif^R$ | Jung et al., 2015 |
| pCCHYC-2 | Integrating plasmid for generation of C-terminal mCherry fusions under control of $P_{Cu}$, $Kan^R$ | Jung et al., 2015 |
| pCCHYC-3 | Integrating plasmid for generation of C-terminal mCherry fusions under control of $P_{Cu}$, $Rif^R$ | Jung et al., 2015 |
| pCCHYN-2 | Integrating plasmid for generation of N-terminal mCherry fusions under control of $P_{Cu}$, $Kan^R$ | Jung et al., 2015 |
| pCVENC-3 | Integrating plasmid for generation of C-terminal Venus fusions under control of $P_{Cu}$, $Rif^R$ | Jung et al., 2015 |
| pdCas9-humanized | Plasmid carrying a codon-optimized version of the *dCas9* gene | Qi et al., 2013 |
| pET21a(+) | Plasmid for overexpression of C-terminally His6-tagged proteins, $Amp^R$ | Novagen |
| pNPTS138 | *sacB*-containing suicide vector used for double homologous recombination, $Kan^R$ | M. R. K. Alley, unpublished |
| pRSFDuet-1 | Plasmid for the coexpression of genes under the control of the T7 promoter | Novagen |
| pRXMCS-2 | Low-copy replicative plasmid for ectopic expression of genes under control of $P_{xyl}$ , $Kan^R$ | Thanbichler et al., 2007 |
| pTB146 | Plasmid for overexpression of N-terminally His6-SUMO-tagged proteins, $Amp^R$ | T. Bernhard (unpublished) |
| pXYFPC-2 | Integrating plasmid for generation of C-terminal eYFP fusions under control of $P_{xyl}$, $Kan^R$ | Thanbichler et al., 2007 |
| pZVENC-2 | Integrating plasmid for generation of C-terminal Venus fusions under control of $P_{Zn}$, $Kan^R$ | Jung et al., 2015 |

##### Table S3. Plasmids generated in this work.

| Plasmid | Description | Construction |
| --- | --- | --- |
| pdCas9Entry | Plasmid carrying (i) humanized *dCas9* under the control of $P_{Cu}$ and (ii) an sgRNA expression cassette comprising the strong constitutive $P_{HNE\_0038}$ promoter followed by a BbsI restriction site, a gene fragment encoding the Cas9 sgRNA handle region and a transcriptional terminator. | a) amplification of a synthetic DNA fragment containing $P_{HNE\_0038}$, a BbsI restriction site, a gene fragment encoding the Cas9 sgRNA handle region and a transcriptional terminator (Integrated DNA Technologies, USA) using primers oJH19 and oJH20<br>b) Gibson assembly of the resulting fragment with NheI-treated pJH01 |
| pEC29 | pNPTS138 derivative for in-frame deletion of *bacD* (*HNE_0444*) | a) amplification of *HNE_0444* upstream and downstream regions from *H. neptunium* chromosomal DNA using the primer pairs HNE_0444_del1/2 and HNE_0444_del3/4<br>b) restriction of the upstream fragment with EcoRI and HindIII and of the downstream fragment with HindIII and NheI<br>c) triple ligation with pNPTS138 cut with EcoRI and NheI |
| pEC32 | pNPTS138 derivative for in-frame deletion of *bacA* (Δ*HNE_2629*) | a) amplification of the *HNE_2629* upstream and downstream regions from *H. neptunium* chromosomal DNA using primer pairs HNE_2629_del1/2 new and HNE_2629_del3/4<br>b) restriction of the upstream fragment with EcoRI and HindIII and of the downstream fragment with EcoRI and NheI<br>c) triple ligation with pNPTS138 cut with HindIII and NheI |
| pEC60 | pCVENC-3 carrying *bacA* | a) amplification of *HNE_2629* from *H. neptunium* chromosomal DNA using primers HNE_2629_for and HNE_2629_comp.rev<br>b) restriction of the PCR product with NdeI and KpnI<br>c) ligation with pCVENC-3 cut with NdeI and KpnI |
| pEC59 | pZVENC-2 harbouring *bacD* | a) amplification of *HNE_0444* from *H. neptunium* chromosomal DNA using primers HNE_0444_for and HNE_0444_rev<br>b) restriction of the PCR product with NdeI and KpnI<br>c) ligation with pZVENC-2 cut with NdeI and KpnI |
| pEC74 | pNPTS138 derivative for the replacement of *bacA* with *bacA-eYFP* | a) amplification of an *HNE_2629-yfp* fragment from pSW56 using primers HNE_2629_HA_for and HNE_2629-FP_eol_rev<br>b) amplification of the *HNE_2629* downstream region from *H. neptunium* chromosomal DNA using primers HNE_2629-FP_eol_for and HNE_2629_del4<br>c) fusion of the two fragments by overlap extension PCR using primers HNE_2629_HA_for and HNE_2629_del4<br>d) restriction of the resulting PCR fragment with HindIII and NheI and ligation with pNPTS138 cut with HindIII and NheI |
| pEC75 | pNPTS138 derivative for the replacement of *bacD* with *bacD-venus* | a) amplification of an *HNE_0444-venus* fragment from pEC59 using primers HNE_0444_for and HNE_0444-FP_eol_rev<br>b) amplification of the *HNE_0444* downstream region from *H. neptunium* chromosomal DNA using primers HNE_0444-FP_eol_for and HNE_0444_integ_rev<br>c) fusion of the two fragments by overlap extension PCR using primers HNE_0444_for2 and HNE_0444_del1<br>d) amplification of a DNA fragment containing *HNE_0444-venus* and the *HNE_0444* downstream region from the resulting PCR product using primers HNE_0444_for3 and HNE_0444_del1extra<br>e) restriction of the PCR fragment with HindIII and NheI and ligation with pNPTS138 cut with HindIII and NheI |
| pEC76 | pNPTS138 derivative for the replacement of *bacD* with *bacD-mCherry* | a) amplification of an *HNE_0444-mCherry* fragment from pEC94 using primers HNE_0444_for2 and HNE_0444-FP_eol_rev<br>b) amplification of the *HNE_0444* downstream region from *H. neptunium* chromosomal DNA using primers HNE_0444FP_eol_for and HNE_0444_del1<br>c) fusion of the two PCR fragments by overlap extension PCR using primers HNE_0444_for3 and HNE_0444_del1<br>d) ) amplification of a DNA fragment containing *HNE_0444-mCherry* and the *HNE_0444* downstream region using primers HNE_0444_for3 and HNE_0444_del1extra<br>e) restriction of the resulting PCR fragment with HindIII and NheI and ligation with pNPTS138 cut with HindIII and NheI |
| pEC86 | pET21a(+) carrying *bacA* | a) amplification of *HNE_2629* from *H. neptunium* chromosomal DNA using primers HNE_2629_for and HNE_2629_rev<br>b) restriction of the PCR product with EcoRI and NdeI<br>c) ligation with pET21a (+) cut with EcoRI and NdeI |
| pEC94 | pCCHYC-3 carrying *bacD* | a) amplification of *HNE_0444* from *H. neptunium* chromosomal DNA using primers HNE_0444_for and HNE_0444_rev<br>b) restriction with of the PCR product with NdeI and KpnI<br>c) ligation with pCCHYC-3 cut with NdeI and KpnI |

#### Table S3. Plasmids generated in this work (continued).

| Plasmid | Description | Construction |
|---|---|---|
| pEC119 | pRSFDuet-1 carrying $P_{T7}$-*bacA-eyfp* | a) amplification of *bacA-yfp* from pEC74 using primers mCherry/venus_rev and NE_2629_for3<br>b) restriction with of the PCR product with PciI and BamHI<br>c) ligation into pRSFDuet-1 cut with NcoI and BamHI |
| pEC120 | pRSFDuet-1 carrying $P_{T7}$-*bacD-ecfp* | a) amplification of *bacD* from *H. neptunium* chromosomal DNA using primers HNE_0444_for and HNE_0444_rev<br>b) restriction of the PCR product with NdeI and KpnI<br>c) ligation into pCCFPC-3 cut with NdeI and KpnI (resulting in pEC70)<br>d) amplification of *bacD-cfp* from pEC70 using primers HNE_0444_for and ecfp_rev2<br>e) restriction of the PCR product with NdeI and MfeI<br>f) ligation into pRSFDuet-1 cut with NdeI and MfeI |
| pEC121 | pRSFDuet-1 carrying $P_{T7}$-*bacA-eyfp* $P_{T7}$-*bacD-ecfp* | a) amplification of *bacB-cfp* using primers HNE_0444_for and ecfp_rev2<br>b) restriction of the PCR product with NdeI and MfeI<br>c) ligation into pEC119 cut with NdeI and MfeI |
| pEC129 | pNPTS138 derivative for the replacement of *rodZ* with *eyfp-rodZ* | a) amplification of *eyfp* from pXYFPC-2 using primers HNE_0620_eol_for2 and HNE_0620_eol_rev2<br>b) amplification of the regions flanking the *eyfp* integration site from *H. neptunium* chromosomal DNA using primer pairs HNE_0620_eol_for/HNE_0620_eol_rev and HNE_0620_eol_for3/HNE_0620_eol_rev3<br>c) fusion of the three PCR fragments by overlap extension PCR using primers HNE_0620_eol_for and HNE_0620_eol_rev3<br>d) restriction of the PCR product with HindIII and NheI and ligation with pNPTS138 cut with HindIII and NheI |
| pJH01 | pCCHYN-2 carrying a codon-optimized version of *dCas9* | a) PCR amplification of *dCas9* from pdCas9-humanized using primers oJH13 and oJH14<br>b) Fusion of the PCR fragment with NdeI/KpnI-treated pCCHYN-2 using Gibson assembly |
| pJH13 | pdCas9Entry carrying an sgRNA targeting *lmdC* | a) phosphorylation and subsequent annealing of oligonucleotides oJH48 and oJH4<br>b) ligation with pdCas9Entry cut with BbsI |
| pMO93 | pNPTS138 derivative for the replacement of *bacA* with $bacA_{F130R}$-*eYFP* | Site-directed mutagenesis of pEC74 with primers bacA-Hn-F130R-for and bacA-Hn-F130R-rev |
| pSP81 | pNPTS138 derivative for in-frame deletion of $lmdC_{Rs}$ (Δ*Rru_A1868*) | a) amplification of the upstream and downstream regions of *Rru_A1868* from *R. rubrum* chromosomal DNA using the primer pairs oSP309/oSP310 and oSP311/oSP312<br>b) insertion of the two fragments into pNPTS138 cut with HindIII and NheI by Gibson assembly |
| pSP82 | pNPTS138 derivative for in-frame deletion of $bacA_{Rs}$ (Δ*Rru_A1867*) | a) amplification of the upstream and downstream regions of *Rru_A1867* from *R. rubrum* chromosomal DNA using the primer pairs oSP315/oSP316 and oSP317/oSP318<br>b) insertion of the two PCR products into pNPTS138 cut with HindIII and NheI by Gibson assembly |
| pSP112 | pRXMCS-2 carrying $P_{lmdC}$-$lmdC_{1\text{-}80}$-*mNeongreen* | a) amplification of $P_{lmdC}$-$lmdC_{(AA1\text{-}80)}$ from *R. rubrum* chromosomal DNA using primers oSP394 and oSP435<br>b) amplification of a fragment encoding a linker and mNeongreen using primers oSP436 and oSP218<br>c) insertion of the two PCR products into pRXMCS-2 cut with NotI and EcoRI by Gibson assembly |
| pSP117 | pNPTS138 derivative for the replacement of *bacA* with *bacA-mCherry* | a) amplification of the regions flanking the *mCherry* integration site from *R. rubrum* chromosomal DNA using the primer pairs oSP321/oSP322 and oSP464/oSP318<br>b) amplification of a fragment encoding a linker and mCherry using primers oSP323 and oSP463<br>c) insertion of the two PCR products into pNPTS138 cut with HindIII and NheI by Gibson assembly |
| pSP118 | pRXMCS-2 carrying $P_{lmdC}$-$bacA_{Rs}$ | a) amplification of $P_{lmdC}$ and *bacA* from *R. rubrum* chromosomal DNA using the primer pairs oSP394/oSP465 and oSP466/oSP467<br>b) insertion of the two PCR products into pRXMCS-2 cut with NotI and EcoRI by Gibson assembly |
| pSP119 | pNPTS138 derivative for the replacement of *bacA* with *bacA-mNeongreen* in the Δ*lmdC* background | a) amplification of the regions flanking the *mNeongreen* integration site from SP68 chromosomal DNA using the primer pairs oSP468/oSP322 and oSP325/oSP318<br>b) amplification of a fragment encoding a linker and mNeongreen using primers oSP323 and oSP324<br>c) insertion of the two PCR products into pNPTS138 cut with HindIII and NheI by Gibson assembly |
| pSP120 | pET21a(+) carrying $lmdC_{Rs}$ | a) amplification of *Rru_A1868* from *R. rubrum* chromosomal DNA using primers oSP469 and oSP470<br>b) insertion of the PCR product into pET21a(+) cut with HindIII and NdeI by Gibson assembly |
| pSP130 | pNPTS138 derivative for in-frame deletion of $lmdC_{Rs}$ (Δ*Rru_A1868*) in the $bacA_{Rs}$::$bacA_{Rs}$-*mCherry* background | a) amplification of the regions flanking *Rru_A1868* from SP109 chromosomal DNA using the primer pairs oSP309/oSP310 and oSP311/oSP485<br>b) insertion of the PCR products into pNPTS138 cut with HindIII and NheI by Gibson assembly |

##### Table S3. Plasmids generated in this work (continued).

| Plasmid | Description | Construction |
|---|---|---|
| pSP198 | pSP112 carrying $P_{lmdC}$-*lmdC*$_{1\text{-}80\ R25A\ H26A\ L27A}$-*mNeongreen* | a) Site-directed mutagenesis of pSP112 with primers oSP690 and oSP691 |
| pSP199 | pSP112 carrying $P_{lmdC}$-*lmdC*$_{1\text{-}80\ R30A\ S31A}$-*mNeongreen* | a) Site-directed mutagenesis of pSP112 with primers oSP692 and oSP693 |
| pSU23 | pNPTS138 derivative for the replacement of *lmdC* with *lmdC*::*lmdC*$_{AA1\text{-}65}$-*HA*-*lmdC*$_{AA66\text{-}405}$ | a) amplification of the regions upstream and downstream of the *HA* integration site from *H. neptunium* chromosomal DNA using the primer pairs oSU36/oSU41 (fragment 1) and oSU39/oSU42 (fragment2)<br>b) amplification of the upstream region from fragment 1 with primers oSU38 and oSU43 (fragment 1.1)<br>c) insertion of fragments 1.1 and 2 into pNPTS138 cut with HindIII and NheI by Gibson assembly |
| pSW56 | pXYFPC-2 carrying *bacA* | a) amplification of *HNE_2629* from *H. neptunium* chromosomal DNA using primers *HNE2629*-for and *HNE2629*-rev<br>b) ligation with pXYFPC-2 cut with EcoRI and NdeI |
| pYL15 | pTB146 carrying *lmdC*$_{226\text{-}345}$ | a) PCR amplification of lmdC with primers lmdC_*M23*_for and *lmdC*_M23_rev<br>b) Insertion of the *lmdC*$_{226\text{-}345}$ fragment into pTB146 cut with BamHI and SapI by Gibson assembly |

**Table S4. PCR primers used in this work.**

| Oligonucleotide | Sequence |
|---|---|
| ecfp_rev2 | tatcaattgttacttgtacagctcgtc |
| HNE_0444_del1 | aaagaattccaggccgaactcgccatcgaaaagg |
| HNE_0444_del2 | tataagcttacagccgtctagttgttctgcagg |
| HNE_0444_del3 | tataagcttatctgctgccatccgcgtgtctccc |
| HNE_0444_del4 | ttttgctagccagttgtgcgcgtgttcgagatcg |
| HNE_0444_for | tatatacatatggcagcagataaggcaagggaaccg |
| HNE_0444_rev | tataggtaccgacggctgtgctggccggcggctc |
| HNE_2629_del1 | tatgaattctcggcggagatcagtccttcatgac |
| HNE_2629_del2 | tataagcttgttattctttgtgaacatgtttgcc |
| HNE_2629_del3 | ttttaagcttccgccgagctgatcgcgcgagggtc |
| HNE_2629_del4 | ttttgctagcacgcgcttgtctgcttcgaggttca |
| HNE_2629_for3 | cgcacatgttcacaaagaataacaaaaccccagc |
| HNE_0444_for | tatatacatatggcagcagataaggcaagggaaccg |
| HNE_0444_rev | tataggtaccgacggctgtgctggccggcggctc |
| HNE_2629_for | ttaacatatgttcacaaagaataacaaaaccccagcggc |
| HNE_2629_rev | tagaattcgagctcggcggcgaggaactcgagatg |
| HNE_2629_comp.rev | tataggtacctcagctcggcggcgaggaactcgag |
| HNE_2629_HA_for | tataagcttatgttcacaaagaataacaaaaccccagc |
| HNE_2629-FP_eol_rev | tcgcgcgatcagctcggcggttacttgtacagctcgtcca |
| HNE_2629-FP_eol_for | tggacgagctgtacaagtaaccgccgagctgatcgcgcga |
| HNE_0444-FP_eol_rev | gtctgcgcaacctgcagaacaattacttgtacagctcgtcca |
| HNE_0444-FP_eol_for | tggacgagctgtacaagtaattgttctgcaggttgcgcagac |
| HNE_0444_for2 | tataagcttatggcagcagataaggcaagggaaccg |
| HNE_0444_for3 | tatagctagcatggcagcagataaggcaagggaaccg |
| HNE_0444_del1extra | tttaagcttcaggccgaactcgccatcgaaaagg |
| HNE_0620_eol_for | tatagctagccgcgctcgaccataaaggt |
| HNE_0620_eol_rev | tcctcgcccttgctcaccatattctaccagtcacttcgac |
| HNE_0620_eol_for2 | gtcgaagtgactggtagaatatggtgagcaagggcgagga |
| HNE_0620_eol_rev2 | tgggtcatgttgtgtgccatatgcatattaattaaggcgc |
| HNE_0620_eol_for3 | gcgccttaattaatatgcatatggcacacaacatgaccca |
| HNE_0620_eol_rev3 | tttaagcttcggccagtgtgcggctgag |
| bacA-Hn-F130R-for | ttcacctggccttcacgcacggcgtttgactgga |
| bacA-Hn-F130R-rev | agtcaaacgccgtgcgtgaaggccaggtgaagcat |
| lmdC_TMH_for | cattcacaggaactcttccatatggcgaagtggagtgcca |
| lmdC_TMH_rev | gccgaccggtgacgcgtaacgttcgcggggccgccgccgacg |
| lmdC_M23_for | gctcacagagaacagattggtggcattcgcgtcgacccct |
| lmdC_M23_rev | gctttgttagcagccggatccttattctttgtgaacatgttt |
| mCherry/venus_rev | ataggatccttacttgtacagctcgtccat |
| oJH13 | acaggaactcttccatatggacaagaagtattctatcggactggccatc |
| oJH14 | cgagatcttaaggtacctcaatccccctccgagctgtgagagg |
| oJH19 | gttacgcgtaccggtggcggccgccatcgtggcgg |
| oJH20 | tcccccgggctgcagctagcaaaaaaagcaccgactcggtgccac |
| oJH48 | accagcgatgatagatctggcgttc |
| oJH49 | aaacgaacgccagatctatcatcgc |
| oSP218 | caccacgtggtacctcgagttacttgtacagctcgtccatgcccatcac |
| oSP309 | gtgcaattgaagccggctggcgccacatgatgccaaagccgccggg |
| oSP310 | catttttccccgctttcaagaaggaaaggtgcagatcctgggggtc |
| oSP311 | ggatctgcacctttccttcttgaaagcggggaaaaatgttttcgaaggc |
| oSP312 | catccggagacgcgtcacggccgaagggccgaccgaatggatcgccc |
| oSP315 | gcaattgaagccggctggcgccaggtgctgctgaccgatttcgatgg |
| oSP316 | cgggttcagggtgccgtcgccggcggtcgaccgtttggagc |
| oSP317 | gctccaaacggtcgaccgccggcgacggcaccctgaaccc |
| oSP318 | catccggagacgcgtcacggccgaagaacaccaagcacaagggggacg |
| oSP321 | gtgcaattgaagccggctggcgccatggccggctgtcgcgc |
| oSP322 | cgcgtaacgttcgaattctccggagctcgggaggccgccgccgg |
| oSP323 | cccggcggcggcctcccgagctccggagaattcgaacgttacg |

**Table S4. PCR primers used in this work (continued)**

| Oligonucleotide | Sequence |
|---|---|
| oSP324 | ccggcagtccaggcggcgattacttgtacagctcgtccatgcccatcac |
| oSP325 | gtgatgggcatggacgagctgtacaagtaatcgccgcctggactgccg |
| oSP394 | cagcgagtcagtgagcgaggaagctcgggtccgggcggcac |
| oSP435 | cctcctcgcccttgctcaccatgatgcgttcgttcttggccgacag |
| oSP436 | cggccaagaacgaacgcatcatggtgagcaagggcgaggaggataac |
| oSP463 | cccggcagtccaggcggcgattacttgtacagctcgtccatgccgc |
| oSP464 | gcggcatggacgagctgtacaagtaatcgccgcctggactgccg |
| oSP465 | gtttggagctagccttcgaaaacatgaggtctcctgttccaaatccgcc |
| oSP466 | gatttggaacaggagacctcatgttttcgaaggctagctccaaacggtc |
| oSP467 | cttaagagctcaccacgtggtacctcgagtcaggaggccgccgccg |
| oSP468 | caattgaagccggctggcgccaatttttttgttgtatcacgtcaaggcg |
| oSP469 | gtttaactttaagaaggagatatacatatgtcggagttcgacccccagg |
| oSP470 | ggtgctcgagtgcggccgcaagcttgccttcgaaaacatttttccccgc |
| oSP485 | ccggagacgcgtcacggccgaagcccacttgaagccctcggggaag |
| oSP690 | gccgccatttccccgatgcggcggcgatggtccgctccgatggg |
| oSP691 | cccatcggagcggaccatcgccgccgcatcggggaaatggcggc |
| oSP692 | cgatcgccacctcatggtcgcggcggatggggcgatgcggc |
| oSP693 | gccgcatcgccccatccgccgcgaccatgaggtggcgatcg |
| oSU36 | cgggccgccgatgttcgtctgt |
| oSU38 | ggagacgcgtcacggccgaagcgggccgccgatgttcgtc |
| oSU39 | ttgaagccggctggcgccacgcggcgaacctttctgatc |
| oSU41 | tacccatacgacgtcccagactacgctcccggcacagcgcagg |
| oSU42 | tctgggacgtcgtatgggtagccgccgccgacgac |
| oSU43 | aaatttcgtcgtcggcggcggctacccatacgacgtcccagact |
